# Integrating the ENCODE blocklist for machine learning quality control of ChIP-seq data with seqQscorer

**DOI:** 10.1101/2025.05.12.653555

**Authors:** Steffen Albrecht, Clarissa Krämer, Philipp Röchner, Johannes U Mayer, Franz Rothlauf, Miguel A Andrade-Navarro, Maximilian Sprang

## Abstract

**Motivation:** Quality assessment of next-generation sequencing data is a complex but important task to ensure correct conclusions from experiments in molecular biology, biomedicine, and biotechnology. We previously introduced seqQscorer, a quality assessment tool using machine learning to support this process. To improve seqQscorer in terms of accuracy and processing time, we integrated the ENCODE blocklist^*^ to derive a new type of quality-related features, supposed to be more informative and faster in generation than those conventionally used by seqQscorer.

**Results:** The novel seqQscorer extension, called seqBLQ, allows us to improve the quality assessment for ChIP-seq data derived from human tissues and cell lines. Furthermore, seqBLQ enhances the usability of the tool by simplifying the installation procedure and reducing the computational resources required for feature generation.

**Availability and implementation:** https://github.com/salbrec/seqQscorer

## Introduction

In this article, we propose *seqBLQ* as an extension for seqQscorer, a machine learning-driven software for automated quality assessment of next-generation sequencing (NGS) samples. Besides whole-genome sequencing, NGS has been coupled with molecular assays to study important drivers of gene regulation and better understand cellular function and cell differentiation. Widely used NGS assays are RNA-seq (RNA sequencing) to quantify gene expression and DNase-seq (DNase I hypersensitive sites sequencing) to detect open chromatin sites (2,3). Open chromatin is a prerequisite for transcription factor binding and is strongly associated with histone modification. As protein-DNA interactions strongly influence gene expression, the ChIP-seq (chromatin immunoprecipitation followed by high-throughput sequencing) assay was developed to reveal such interactions (4). These assays are essential for both fundamental research and biomedical studies investigating cancer and other life-threatening diseases (5–8).

Technological advancements for NGS allowed scientists to create an enormous amount of data, filling publicly available databases (9–11). These advancements also reduced costs, resulting in research projects with increasing numbers of samples (12). While these large NGS datasets collectively allow for a multiplicity of analyses, a key challenge that remains is sample quality assessment. Low-quality samples must be accurately identified to be refined or even replaced to ensure reliable downstream analyses. This is even more important when sequencing results are involved in decision-making processes for precision medicine (13).

While quality assessment is an important task, it is challenging and requires the experience of domain experts. Several tools exist to support this process, providing quality reports incorporating different statistics derived from NGS samples. Some of these are generic to different assays (14,15), while others are tailored to specific molecular assays (16,17). These reports can be comprehensive, providing an overview of the quality of NGS samples from multiple perspectives (14,18). However, the inspection of this information can be time-consuming and requires domain expertise (19).

To reduce manual work, we previously introduced *seqQscorer*, which applies machine learning (ML) models trained on quality-related reports and statistics to automate the quality assessment of NGS samples from RNA-seq, ChIP-seq, DNase-seq, and ATAC-seq (an alternative to DNase-seq). The models are trained by supervised ML algorithms and return a single quality score that is used to rank potentially large sets of NGS samples by their quality. The quality labels used for model training are defined by the manually curated ENCODE status, considering *revoked* fastq files as low-quality and *released* fastq files as high-quality NGS samples.

In the original and following work, we showed that the seqQscorer ML models capture quality issues across assays and species and are generalizable to data from other public databases (19,20). Additionally, its quality score relates to batch effects (21), and it enabled us to introduce *Quality Imbalances*, a new type of technical bias that arises in experiments dealing with multiple samples (e.g., controls and conditions, with several replicates) when the mean quality scores of the groups of interest (e.g., control and treatment) strongly differ (22). While seqQscorer achieves remarkable results, using commonly used quality-related features derived from sequencing samples, we continued to further explore possibilities to improve seqQscorer in terms of usability, processing time, and accuracy. While the accuracy of the conventional seqQscorer is near-perfect for some subsets, such as the one for RNA-seq, there are areas for improvement, especially for some ChIP-seq data subsets.

This led us to investigate the ENCODE blocklist, a set of problematic genomic regions defined based on data from the ENCODE portal (1). These regions can, for instance, contain collapsed repeats in the genome assembly that lead to a high number of mapped reads, in this case considered as an artifact in the sequencing signal. It has been shown that removing these regions prior to downstream analysis is important to achieve reliable results, and bioinformatic analysis tools incorporate automated filtering of these regions, thereby removing artifacts before downstream analysis (23). According to the original work, the blocklisted regions can also be used to compute a single-value quality metric describing the fraction of reads that overlap with any of these regions.

Given the strong relationship between these blocklisted regions and data quality, we here explore the potential of creating ML features based on these regions to train quality assessment models as introduced by seqQscorer. In comparison to a single value for the fraction of reads overlapping with any blocklist region, the *blocklist features* we propose and integrate into seqBLQ are described by multiple values describing the number of overlapping reads for each blocklisted region. We show that seqBLQ performs well on data across assays and organisms and can even improve model performance for ChIP-seq data derived from human tissues and cell lines.

## Datasets and Methods

The datasets used in the following experiments are the same as those used in the seqQscorer study, namely, samples derived from tissues and cell lines, called *biosamples*, from two species, human and mouse. Sequencing samples have been obtained by the ENCODE consortium using three different assays (ChIP-seq, DNase-seq, RNA-seq), further specified by two sequencing modes (single-end, paired-end). Importantly, the samples have been labelled using the ENCODE status, an attribute assigned to experiments and single samples, manually curated by the Data Coordination Center (DCC). Files with the status *revoked* receive the low-quality label, while *released* samples represent high-quality data. Bioinformatics tools have been applied to derive comprehensive quality-related information from the raw sequencing samples to assemble the conventional feature sets. For more details, we refer to the seqQscorer study (19).

To create the blocklist features for seqBLQ, we first mapped the sequencing reads for each raw sequencing sample, provided in *fastq* format. As done for seqQscorer, we used *Bowtie2* with default settings consistently across samples to align sequencing reads against the genome assemblies *hg38* and *mm10* for human and mouse, respectively. Then, for each blocklisted region, a feature value was created by counting the overlapping mapped reads. According to the size of the blocklists, 636 regions for human and 3435 regions for mouse, this results in 636 and 3435 features for human and mouse data, respectively. Additionally, we created blocklist features using reads mapped against smaller genome assemblies restricted to the genomic regions from the blocklists (**Fig. 1A**). The latter has been done to verify that using a blocklist-restricted mapping has no negative impact on the accuracy of the quality assessment, which would allow us to simplify the usage of the tool by using restricted reference genomes for the mapping.

**Figure 1.**
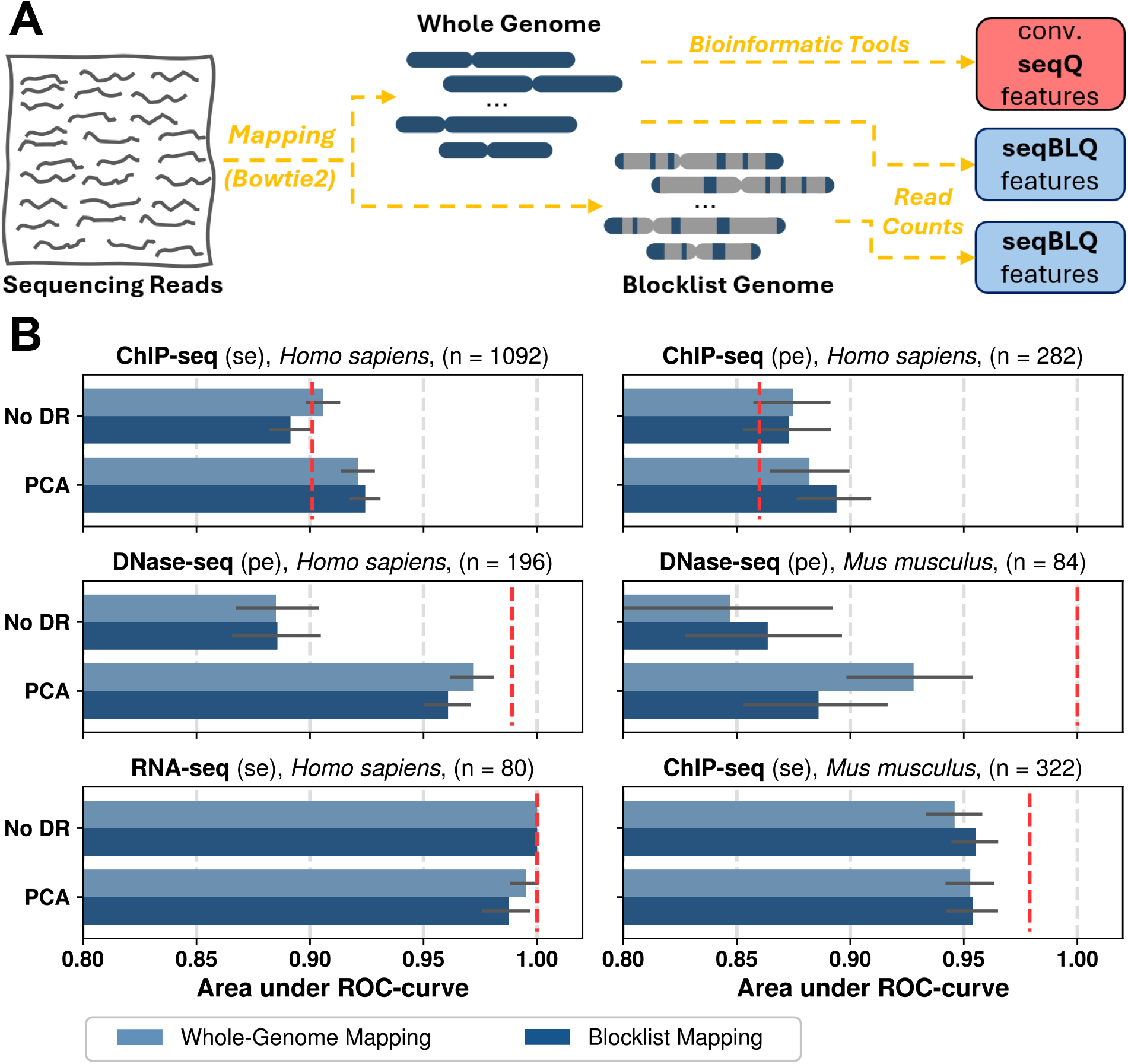
Data preprocessing and performance of the seqBLQ quality assessment. **A** Simplified data flow for the conventional seqQ feature engineering using the whole-genome mapping and the seqBLQ feature engineering that can be performed with a blocklist-restricted genome assembly. **B** seqBLQ model performance using the Random Forest algorithm with default settings, see the scikit-learn package (27). The bars represent the mean area under the ROC curve for these models evaluated on multiple training/testing splits. The error bars show the 95% confidence intervals of the performance measures obtained. Subtitles summarize the characteristics of the underlying datasets with assay name, organism, and run type, with (se) and (pe) abbreviating single-end and paired-end, respectively. The number “n” refers to the number of samples in the subsets. The dashed red line shows the mean performance of models that were fine-tuned on the specific subsets during a comprehensive ML analysis performed for the original seqQscorer study. Note that a few samples had been removed from the ENCODE portal in the meantime, resulting in slightly lower sample sizes for some subsets compared to the subsets originally used.

The ML experiments aim to compare the quality classification performance of ML models trained with the conventional seqQscorer features (seqQ) and the new blocklist features (seqBLQ). Models were evaluated by the area under the ROC curve (auROC) within a ten-fold cross-validation. To obtain a statistically robust model evaluation, we repeated the cross-validation five times to generate 50 training-testing splits.

As the number of blocklist features used in seqBLQ is high, we face the challenge of small data learning that occurs for datasets with a high number of ML features but a comparatively low number of samples (24,25). Therefore, we integrate dimensionality reduction using Principal Component Analysis (PCA) (26). Fitted on the training set, PCA is used to transform training and testing sets to reduce the dimensionality, keeping principal components that explain most of the data variance. The number of relevant components is automatically determined by adding components until 99.99% of the training data variance is explained.

## Results

For ChIP-seq data, derived from human biosamples, we observed a clear improvement in model performance achieved by seqBLQ (**Fig. 1B**). For this data, the accuracy can be further improved by using the blocklist-restricted read mapping, especially in combination with the PCA dimensionality reduction that, as expected, proves to be necessary in general to maintain high model performance. Depending on the data subsets and feature sets used, the number of principal components used differs (**Table S1**). For data from mouse biosamples or assays different from ChIP-seq, seqBLQ achieves high area under the ROC curve scores. However, the highest model performance is achieved by the conventional seqQscorer approach (**Fig. 1B**).

## Discussion and Outlook

Our results clearly advise users of seqQscorer to apply the seqBLQ extension when assessing the quality of ChIP-seq data from human biosamples.

The ENCODE blocklists have been derived from a collection of control ChIP-seq samples, which are randomly sheared DNA regions from non-immunoprecipitated chromatin, also called *ChIP-seq input*. It is remarkable that seqBLQ achieves high assessment performance, also for data from DNase-seq and RNA-seq. This indicates that quality-related information from problematic regions identified through ChIP-seq input samples can be transferred to samples from DNase-seq and RNA-seq. For these assays, however, the best quality assessment is still achieved by the conventional seqQscorer approach.

Additionally, the conventional seqQscorer achieves very high performance for ChIP-seq data from mouse biosamples, and the proposed blocklist features used in seqBLQ could not be used to perform at the same level. A possible explanation for this could be that the mouse blocklist is not as reliable in pointing to problematic regions as the human blocklist, due to lower data availability of ChIP-seq input samples from mouse biosamples used to create the blocklist (1).

The blocklists are already used to create a single-value quality control metric described by *the fraction of reads in blocklisted regions* (1), here called *FRiBL*. As the FRiBL has been considered by the ENCODE DCC when samples got revoked in the past, a statistical relationship could potentially exist between the blocklist features we propose and the ENCODE status used to label the data. However, we assume that the FRiBL, as a single-value quality control metric, does not capture the level of detail encoded in the blocklist features we propose, represented by a count of mapped reads for each of these regions. Furthermore, we expect that the FRiBL metric is not always the sole reason for sample revocation, and that the seqBLQ quality score relates to a wider range of quality-related problems. Indeed, the FRiBL is poorly related to the quality labels, while the seqBLQ quality score allows for a clear differentiation between low- and high-quality samples (**Fig. S1**).

Besides the improved model performance that can be achieved by using the blocklist feature extension of seqQscorer, there are other advantages of the proposed extension. As the blocklist features can be derived from mapped reads using a blocklist-restricted genome index, the mapping is about twice as fast (runtime comparisons not presented). More importantly, the blocklist-restricted genome index files are small, which allows us to provide the ready-to-use Bowtie2 index files within the GitHub repository. Finally, the blocklist feature engineering requires fewer bioinformatics tools, which simplifies the installation of dependencies.

## Code and Data Availability

The new blocklist feature engineering and seqBLQ quality scoring have been added to the seqQscorer GitHub repository: https://github.com/salbrec/seqQscorer

This repository is maintained and updated regularly by SA and MS. The datasets and blocklist-restricted genome index files used in this study can also be downloaded from there.

## Acknowledgements

The authors gratefully acknowledge the computing time provided to them at the NHR Center NHR@SW at Johannes Gutenberg University Mainz. This is funded by the Federal Ministry of Education and Research, and the state governments participating on the basis of the resolutions of the GWK for national high-performance computing at universities (www.nhr-verein.de/unsere-partner).

## References

1. Amemiya HM, Kundaje A, Boyle AP. The ENCODE blacklist: identification of problematic regions of the genome. Scientific reports. 2019;9(1):9354.

2. Ozsolak F, Milos PM. RNA sequencing: advances, challenges and opportunities. Nature reviews genetics. 2011;12(2):87–98.

3. Song L, Crawford GE. DNase-seq: a high-resolution technique for mapping active gene regulatory elements across the genome from mammalian cells. Cold Spring Harbor Protocols. 2010;2010(2):pdb–prot5384.

4. Mardis ER. ChIP-seq: welcome to the new frontier. Nature methods. 2007;4(8):613–4.

5. Wang C, Han B. Twenty years of rice genomics research: From sequencing and functional genomics to quantitative genomics. Molecular Plant. 2022;15(4):593–619.

6. Satam H, Joshi K, Mangrolia U, Waghoo S, Zaidi G, Rawool S, et al. Next-generation sequencing technology: current trends and advancements. Biology. 2023;12(7):997.

7. Prasher D, Greenway SC, Singh RB. The impact of epigenetics on cardiovascular disease. Biochem Cell Biol. 2020 Feb;98(1):12–22.

8. Esteller M. Epigenetics in Cancer. N Engl J Med. 2008 Mar 13;358(11):1148–59.

9. de Souza N. The ENCODE project. Nature methods. 2012;9(11):1046–1046.

10. Zheng R, Wan C, Mei S, Qin Q, Wu Q, Sun H, et al. Cistrome Data Browser: expanded datasets and new tools for gene regulatory analysis. Nucleic acids research. 2019;47(D1):D729–35.

11. Clough E, Barrett T. The gene expression omnibus database. In: Statistical genomics. Springer; 2016. p. 93–110.

12. Kumar KR, Cowley MJ, Davis RL. Next-Generation Sequencing and Emerging Technologies*. Semin Thromb Hemost. 2024 Oct;50(07):1026–38.

13. Mosele MF, Westphalen CB, Stenzinger A, Barlesi F, Bayle A, Bièche I, et al. Recommendations for the use of next-generation sequencing (NGS) for patients with advanced cancer in 2024: a report from the ESMO Precision Medicine Working Group. Annals of Oncology. 2024;35(7):588– 606.

14. Brown J, Pirrung M, McCue LA. FQC Dashboard: integrates FastQC results into a web-based, interactive, and extensible FASTQ quality control tool. Bioinformatics. 2017;33(19):3137–9.

15. Meyer CA, Liu XS. Identifying and mitigating bias in next-generation sequencing methods for chromatin biology. Nature Reviews Genetics. 2014;15(11):709–21.

16. D. Chikina M, G. Troyanskaya O. An effective statistical evaluation of ChIPseq dataset similarity. Bioinformatics. 2012;28(5):607–13.

17. Trapnell C, Pachter L, Salzberg SL. TopHat: discovering splice junctions with RNA-Seq. Bioinformatics. 2009;25(9):1105–11.

18. Ewels P, Magnusson M, Lundin S, Käller M. MultiQC: summarize analysis results for multiple tools and samples in a single report. Bioinformatics. 2016;32(19):3047–8.

19. Albrecht S, Sprang M, Andrade-Navarro MA, Fontaine JF. seqQscorer: automated quality control of next-generation sequencing data using machine learning. Genome biology. 2021;22(1):1–20.

20. Sprang M, Krüger M, Andrade-Navarro MA, Fontaine JF. Statistical guidelines for quality control of next-generation sequencing techniques. Life Science Alliance [Internet]. 2021 [cited 2025 May 10];4(11). Available from: https://www.life-science-alliance.org/content/4/11/e202101113.abstract

21. Sprang M, Andrade-Navarro MA, Fontaine JF. Batch effect detection and correction in RNA-seq data using machine-learning-based automated assessment of quality. BMC bioinformatics. 2022;23(6):1–15.

22. Sprang M, Möllmann J, Andrade-Navarro MA, Fontaine JF. Overlooked poor-quality patient samples in sequencing data impair reproducibility of published clinically relevant datasets. Genome Biol. 2024 Aug 16;25(1):222.

23. Stark R, Brown G. DiffBind: differential binding analysis of ChIP-Seq peak data. R package version. 2011;100(4.3).

24. Vecchi E, Pospíšil L, Albrecht S, O’Kane TJ, Horenko I. eSPA+: Scalable entropy-optimal machine learning classification for small data problems. Neural Computation. 2022;34(5):1220–55.

25. Vecchi E, Berra G, Albrecht S, Gagliardini P, Horenko I. Entropic approximate learning for financial decision-making in the small data regime. Research in International Business and Finance. 2023;65:101958.

26. Kherif F, Latypova A. Principal component analysis. In: Machine learning [Internet]. Elsevier; 2020 [cited 2025 May 10]. p. 209–25. Available from: https://www.sciencedirect.com/science/article/pii/B9780128157398000122

27. Pedregosa F, Varoquaux G, Gramfort A, Michel V, Thirion B, Grisel O, et al. Scikit-learn: Machine learning in Python. the Journal of machine Learning research. 2011;12:2825–30.

