## Supplementary material for "Integrating the ENCODE blocklist for machine learning quality control of ChIP-seq data with seqQscorer": Table S1, Fig. S1

---

* Following inclusive language recommendations, we use the term ENCODE blocklist, previously introduced as the ENCODE blacklist

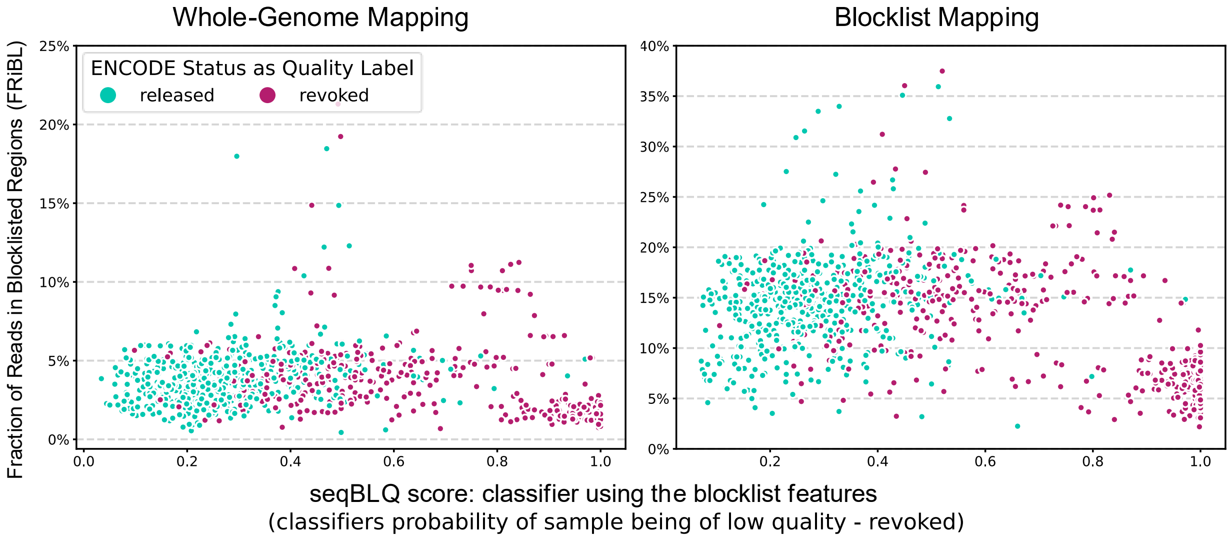


**Figure S1 – Comparing the proposed seqBLQ quality score with the FRiBL quality metric**

*Dots in the scatter plots represent the 1092 samples from the single-end ChIP-seq subset for human biosamples. The score on the y-axis shows the fraction of reads in blocklisted regions (FRiBL). The score on the x-axis is the score from the seqQscorer model, making use of the more complex blocklist features. The higher this score, the lower the expected quality, as these scores are essentially the probability of a sample being of low quality. The color of the dots describes the quality label derived from the ENCODE status, using revoked samples as low-quality and released samples as high-quality data.*

*Especially for the low-quality samples, visible in the lower-right corner in both panels, the seqBLQ score clearly identifies those as low quality through the high classifier probabilities. The FRiBL does not vary strongly between samples in general, and there is no clear trend for low-quality samples to have a higher FRiBL.*

*Note that the FRiBL is lower when using the full genome assembly, as multi-mapped reads can randomly be assigned to non-blocklist regions when the full genome is available.*

| | | | Number of PCs explaining 99.99% of the Variance | | |
|---|---|---|---|---|---|
| **Dataset Specification** | ***n*** **samples, *n* features** | **Mapping** | **min** | **median** | **max** |
| **ChIP-seq (se), human** | 1092, 636 | Whole Genome | 75 | 76 | 81 |
| | | Blocklist-Restricted | 74 | 76 | 78 |
| **ChIP-seq (pe), human** | 282, 636 | Whole Genome | 56 | 57 | 60 |
| | | Blocklist-Restricted | 59 | 61 | 63 |
| **DNase-seq (pe), human** | 196, 636 | Whole Genome | 21 | 23 | 25 |
| | | Blocklist-Restricted | 15 | 17 | 19 |
| **DNase-seq (pe), mouse** | 84, 3435 | Whole Genome | 9 | 11 | 12 |
| | | Blocklist-Restricted | 10 | 11 | 12 |
| **RNA-seq (se), human** | 80, 636 | Whole Genome | 3 | 3 | 3 |
| | | Blocklist-Restricted | 4 | 5 | 5 |
| **ChIP-seq (se), mouse** | 322, 3435 | Whole Genome | 12 | 12 | 14 |
| | | Blocklist-Restricted | 13 | 14 | 15 |

**Table S1 – Data dimensionality after dimensionality reduction using the PCA**

*The PCA is fitted based on the training set for each training-testing split. There are 50 splits, and as the PCA is fitted on the training data, the number of principal components (PCs) slightly varies between the training-testing splits. This table provides an overview of the number of PCs used by the classifiers after dimensionality reduction. Across the 50 splits, we here report the minimum (min), maximum (max), and median number of PCs required to explain 99.99% of the variance of the training data. The full blocklist feature sets are size 636 and 3435 for human and mouse, respectively.*
